# The SWIB domain-containing DNA topoisomerase I of *Chlamydia trachomatis* mediates DNA relaxation

**DOI:** 10.1101/2024.12.03.626651

**Authors:** Li Shen, Caitlynn Diggs, Shomita Ferdous, Amanda Santos, Neol Wolf, Andrew Terrebonne, Luis Lorenzo Carvajal, Guangming Zhong, Scot P. Ouellette, Yuk-Ching Tse-Dinh

## Abstract

The obligate intracellular bacterial pathogen, *Chlamydia trachomatis* (Ct), has a distinct DNA topoisomerase I (TopA) with a C-terminal domain (CTD) homologous to eukaryotic SWIB domains. Despite the lack of sequence similarity at the CTDs between *C. trachomatis* TopA (CtTopA) and *Escherichia coli* TopA (EcTopA), full-length CtTopA removed negative DNA supercoils *in vitro* and complemented the growth defect of an *E. coli topA* mutant. We demonstrated that CtTopA is less processive in DNA relaxation than EcTopA in dose-response and time course studies. An antibody generated against the SWIB domain of CtTopA specifically recognized CtTopA but not EcTopA or *Mycobacterium tuberculosis* TopA (MtTopA), consistent with the sequence differences in their CTDs. The endogenous CtTopA protein is expressed at a relatively high level during the middle and late developmental stages of *C. trachomatis*. Conditional knockdown of *topA* expression using CRISPRi in *C. trachomatis* resulted in not only a developmental defect but also in the downregulation of genes linked to nucleotide acquisition from the host cells. Because SWIB-containing proteins are not found in prokaryotes beyond *Chlamydia* spp., these results imply a significant function for the SWIB-containing CtTopA in facilitating the energy metabolism of *C. trachomatis* for its unique intracellular growth.

**Importance:** *C. trachomatis* (Ct) is a medically important bacterial pathogen that is responsible for the most prevalent sexually transmitted bacterial infection. Bioinformatics, genetics, and biochemical analyses have established that the presence of a SWIB domain in CtTopA, a DNA topoisomerase I, is relevant to chlamydial physiology. Further defining the mechanisms of the C-terminal SWIB domain on the catalytic function of CtTopA in an intracellular pathogen is warranted for a more complete understanding of the interactions between *C. trachomatis* and its host cells.

## Introduction

DNA topoisomerases (Topos) are essential enzymes maintaining DNA supercoiling at appropriate levels in all live cells (1–3). Depending on their actions on DNA, Topos can be broadly divided into type I and type II. Type I Topos (e.g. TopA or TopoI) transiently cleave and reseal a single strand of the DNA helix in the absence of ATP. Type II Topos (e.g., DNA gyrase and TopoIV) cut and religate both DNA strands in the presence of ATP. In *Escherichia coli*, DNA supercoiling is chiefly balanced by the contrasting functions of DNA-relaxing TopA and the negative supercoiling-introducing DNA gyrase. The main function of TopoIV is to disentangle replicated DNA enabling the segregation of duplicated chromosomes. Bacterial type II Topos are targets for fluoroquinolone, a class of clinically relevant antibiotics. With the alarming rise of antibiotic resistance, great efforts have been given to develop novel poison or catalytic inhibitors of Topos for use against difficult-to-treat bacterial infections (4, 5), including drug-resistant *Mycobacteria tuberculosis* and *Neisseria gonorrhoeae*.

*Chlamydia trachomatis* is the leading cause of bacterial sexually transmitted infections (STI) (6, 7). In 2020, an estimated 128.5 million new *C. trachomatis* cases occurred worldwide among individuals aged 15 to 49 years (8, 9). Over 50% of men and >75% of women with *C. trachomatis* infection are asymptomatic. The lack of durable immunity in most individuals can result in recurrent or chronic *C. trachomatis* infection. In women, this can lead to pelvic inflammatory disease and eventually to ectopic pregnancy and tubal factor infertility. *C. trachomatis* can be transmitted to newborns during vaginal birth, causing conjunctivitis and pneumonia. *C. trachomatis* infections have also been epidemiologically associated with gynecologic cancers and a greater risk of acquiring HIV or other STIs. Co-infections of *C. trachomatis* and other STI pathogens, such as *N. gonorrhoeae,* are common. Although *C. trachomatis* infection can be cured by antibiotics, a compelling whole genome sequence study indicated that *C. trachomatis* can establish chronic infections even with repeated antibiotic treatments, but the reasons are unknown (10). There is an urgent need for improved preventative and treatment strategies to solve the problems associated with *C. trachomatis* infection.

As an obligate intracellular bacterium, *C. trachomatis* lives within a membrane vacuole (inclusion) and relies on host energy and nutrient resources (11, 12). *C. trachomatis* undergoes a characteristic developmental cycle involving morphologically and functionally divergent forms that differentiate between them at early and late stages of the cycle. The elementary body (EB) is a small, infectious, metabolically limited form with highly condensed chromatin. The reticulate body (RB) is a replicative and more metabolically active form with dispersed chromatin. The observations that the *C. trachomatis* developmental cycle correlates to temporal gene expression (13, 14) and differential plasmid DNA supercoiling levels (15–17) led to the assumption that DNA supercoiling is a global regulator provoking chlamydial developmental changes. This notion is supported by *in vitro* studies showing that selected early and midcycle promoters of *C. trachomatis* are sensitive to changing DNA supercoiling levels (17–19). We recently utilized CRISPRi technology for conditional repression of *C. trachomatis topA* encoding TopA (CtTopA) to bypass lethality issues associated with the disruption of essential genes (20). Our results have demonstrated that targeted knockdown of *topA* in *C. trachomatis* impairs RB-to-EB transition, leads to downregulation of EB-associated gene expression, and results in a greater sensitivity of *C. trachomatis* to the fluoroquinolone moxifloxacin. Repression of *topA* also affected gyrase expression, indicating a potential compensatory mechanism for survival to offset TopA deficiency. These data highlight the importance of CtTopA in the chlamydial developmental cycle.

It remains unknown how CtTopA acts to affect *C. trachomatis* physiology. Since 1998, when the first *C. trachomatis* genome was published, it has been predicted that a eukaryotic SWIB domain is fused to the C-terminus of the canonical conserved catalytic domains of TopA (21); however, no study was performed to understand its significance. The SWIB-containing protein is widely present in eukaryotes but only rarely found in prokaryotes, except for *Chlamydia* spp. This prompted us to hypothesize that the SWIB-containing CtTopA has a critical function in *Chlamydia* biology – specifically, the chlamydial developmental cycle. Here, we characterized the SWIB-domain containing CtTopA by (i) determining DNA relaxation capacity of CtTopA compared to that of well-studied EcTopA *in vitro* and in *E. coli topA* mutant strains, and (ii) assessing CtTopA’s expression levels and the impact on *C. trachomatis* nucleotide metabolism in the context of infection. Our studies provide strong evidence that *C. trachomatis* naturally produces and operates a functional SWIB-containing CtTopA that participates in the regulation of the chlamydial developmental cycle and nucleotide metabolism.

## Results

### The C-terminal SWIB domain of TopA is unique to *Chlamydia* spp - bioinformatics evidence

Bacterial Topo I proteins consist of two critical functional regions: conserved N-terminal domains (NTDs), which have DNA cleavage and religation activities, and highly diverse C-terminal domains (CTDs), which are crucial for the ability to relax DNA and to exert other catalytic activities (1–3). Two prototypes of CTD motifs, Topo_C_ZnRpt and Topo_C_Rpt, were originally identified in EcTopA (22, 23) and *M. tuberculosis* TopA (MtTopA) (24), respectively. In addition, a CTD tail of lysine repeats was mainly associated with TopAs from *M. tuberculosis* and other GC-rich Actinobacteria phylum members (25). *C. trachomatis* has evolved to have a small, AT-rich chromosome (21). We initially analyzed the domain composition of CtTopA using InterPro (26). Its amino acid sequences were then specifically aligned with those of TopA counterparts in medically important bacteria, *E. coli, M. tuberculosis*, *Helicobacter pylori*, *Pseudomonas aeruginosa*, and *N. gonorrhoeae.* As shown in Fig S1 and Fig.1a-c, the NTDs of TopAs (corresponding to EcTopA D1-D4) contain conserved domains, including topoisomerase-primase domain (TOPRIM) and DNA-binding sites (6,7). However, the CTDs (corresponding to EcTopA D5-D9) are varied. The EcTopA contains three 4-cysteine (4C) zinc finger motifs (D5–D7) and two zinc ribbon-like motifs (D8–D9) at its CTD. The 4C zinc fingers are also present in the TopAs from *C. trachomatis* (three), *P. aeruginosa* (three), *H. pylori* (four), and *N. gonorrhoeae* (four), but not in *M. tuberculosis* TopA, which instead has 4 Topo_C_Rpt domains and 2 lysine repeats. Specifically, CtTopA stands out for possessing a eukaryotic SWIB-domain at its far CTD (in the place of EcTopA zinc ribbon-like domains D8-D9). AlphaFold prediction determined that these amino acid sequences folded into a SWIB-like three-dimensional shape (Fig.1d) distinct from other known structures of TopA proteins (2, 27), suggesting a potentially novel protein fold or functional variation in the TopA family. The Basic Local Alignment Search Tool (BLASTp) found that the TopA from members of the Chlamydiaceae family are conserved in harboring the SWIB domain. Table S1 shows alignment of amino acid residues from the top 500 homologues of CtTopA (corresponding to amino acids 750-857) in *Chlamydia* spp. Thus, CtTopA is distinguished from other bacterial TopAs by its unusual SWIB domain at its CTD, suggesting potentially unique functions for this TopA ortholog.

### Recombinant CtTopA is enzymatically active in DNA relaxation *in vitro*

We sought to directly determine the DNA relaxation activity of CtTopA *in vitro*. Thus, we created a plasmid encoding CtTopA protein with an N-terminal 6xHis tag under the control of the T7 promoter and transformed it into *E. coli* BL21(DE3). Protein expression was induced by adding isopropyl β-D-1-thiogalactopyranoside (IPTG). The recombinant CtTopA was purified to homogeneity (Fig. 2a) and used for DNA relaxation assays by comparing its activity to that of EcTopA. With serial dilutions of CtTopA or EcTopA (at concentrations from 0 to 50 nM) and constant amounts of negatively supercoiled plasmid DNA, we observed different patterns of DNA relaxation. More CtTopA protein is required for the DNA substrate to reach a fully relaxed state (Fig. 2b) with some supercoiled DNA substrate remaining at low enzyme levels (1.5 nM or less) at the end of 30 min incubation. Additionally, a time-course study was performed with incubation of 25 nM of CtTopA or EcTopA and constant amounts of plasmid DNA for 0-30 minutes. The reaction products from CtTopA relaxation have fewer bands in the gel corresponding to the entire population of plasmids having similar number of superhelical turns removed during the time course of relaxation (Fig. 2c-d). Longer incubation is required for the DNA substrate to reach a fully relaxed state as reflected by measurement of the percentage of DNA relaxation (Figs. 2d and S2). In contrast, the EcTopA relaxation is more processive, with the enzyme staying bound to the plasmid substrate to remove nearly all the superhelical turns instead of dissociating from the DNA substrate after removing only a few superhelical turns. Collectively, these results imply that CtTopA is less efficient than EcTopA in relaxation of negatively supercoiled DNA.

### CtTopA expression complements *E. coli* strains with *topA* mutations

Next, we examined the activity of CtTopA in the *E. coli topA* mutant strains (Table S2). *E. coli* VS111-K2 (28) is cold sensitive and has a growth defect at 30°C due to the *ΔtopA* mutation resulting in excessive negative DNA supercoiling. If CtTopA functions in *E. coli*, then bacterial growth should be improved when it is expressed at 30°C. We transformed a pBOMB-based shuttle plasmid expressing P*_tet_*-controlled *C. trachomatis topA* (20) or not into VS111-K2. After incubation on LB agar plates at 30°C for 18 h, the CtTopA expressing strain exhibited better growth than the vector control strain regardless of the addition of inducer anhydrotetracycline (aTC) (Fig. 3a), suggesting leaky expression of CtTopA in uninducing conditions. Both strains grew well at 37°C, as expected, and did not grow at 42°C (data not shown). The latter might be due to the influence of the chlamydial plasmid encoded gene products (e.g., 8 open reading frames) (29). The capacity of CtTopA to complement was further supported by a growth curve assay (Fig. 3b). The expression of CtTopA was confirmed by immunoblotting (Fig. 3b), indicating CtTopA was expressed in the absence and presence of aTC in *E. coli*.

To recapitulate and to evaluate the complementing efficiency, we used *E. coli* strain AS17 (30–32). The *topA* gene in AS17 has a G65N mutation and an amber codon instead of the W79 residue found in wild-type EcTopA. Studies have shown that AS17 is not viable for growth at 42°C because of lack of relaxation activity from the chromosomally encoded EcTopA at the non-permissive temperature (30, 31). However, background noninduced expression of bacterial Topo I under the control of the T7 promoter in an expression plasmid can complement growth of *E. coli* AS17 at 42°C (31). We transformed the pET-based plasmid expressing *C. trachomatis topA* or *E. coli topA* under the control of the T7 promoter into AS17. The empty vector containing strain was used as the control. We observed that the CtTopA expressing clone supported growth of *E. coli* AS17 at 42°C (Fig. 3c). Compared to the EcTopA-expressing positive control, the CtTopA clone grew at about 10-fold lower efficiency, consistent with the less robust relaxation activity for CtTopA in the *in vitro* enzyme activity assay (Fig. 2b-d). Nevertheless, these results indicate that expression of basal levels of CtTopA is necessary and sufficient to correct the growth defect of *E. coli topA* mutants.

**Figure 1.**
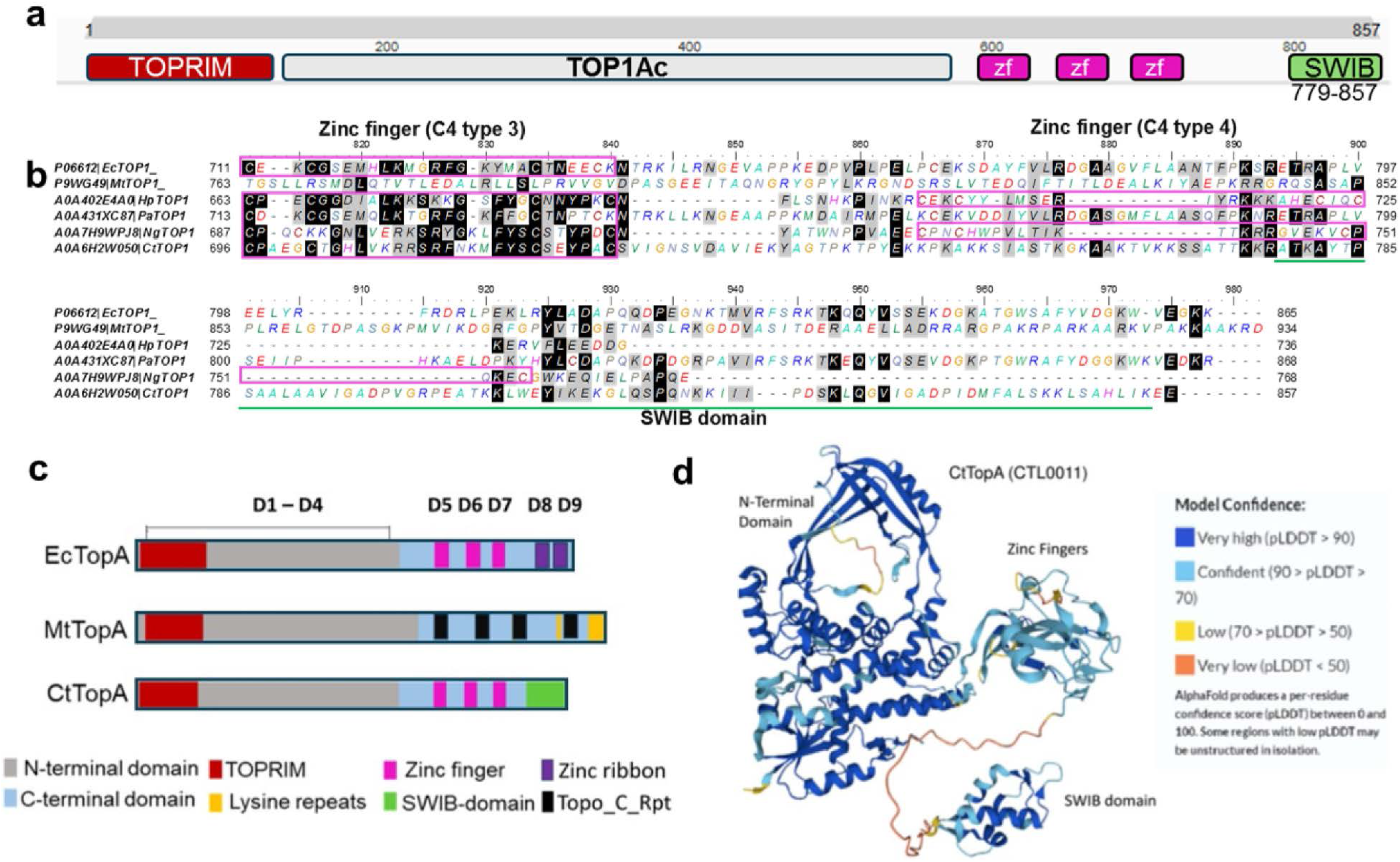
C-terminal SWIB domain is unique in CtTopA. **(a)** Domain composition of CtTopA (CTL0011) predicted by InterPro. Zf: 4C zinc fingers. **(b)** Alignment of amino acid residues of the CTDs of TopAs from *E. coli, M. tuberculosis*, *H. pylori*, *P. aeruginosa*, *N. gonorrhoeae* and *C. trachomatis*. Accession numbers are shown on the left. The conserved 4C zinc fingers are boxed. The position of SWIB-domain in CtTopA is underlined (green). ClustalW was used for alignment with Matric BLOSUM62. See Fig. S1 for entire sequence alignments of these bacterial TopAs. (**c**) Schematic diagram showing domains of the EcTopA (D1-D9) compared to domains found in MtTopA and CtTopA. The gray or light blue bar represents the N- and C-terminal domains. The TOPRIM (red), zinc finger (cyan), Topo_C-Rpt (black), lysine repeats (yellow), and SWIB domain (green) are as indicated. (**c**) Structural model of CtTopA by AlphaFold. The NTD, CTD zinc fingers, and the SWIB domain are as indicated. Model confidences are shown on right.

**Figure 2.**
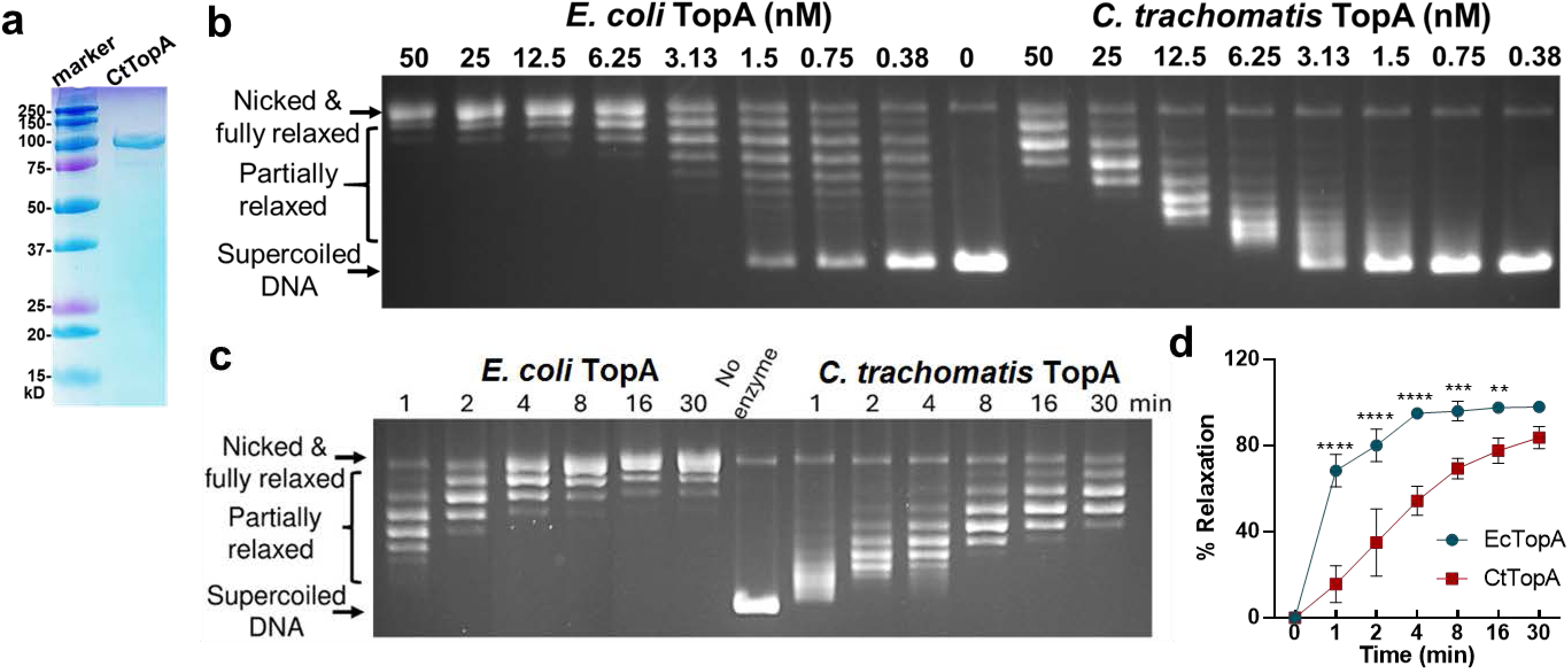
Comparison of the *in vitro* DNA relaxation activity of recombinant CtTopA to EcTopA. (**a**) SDS-PAGE/coomassie staining gel showing recombinant CtTopA protein purified from *E. coli.* (**b**) Concentration-dependent DNA relaxation. Serial dilutions of EcTopA and CtTopA as indicated were incubated with 0.3 μg (5.2 nM) negatively supercoiled DNA for 30 min, followed by agarose gel electrophoresis. (**c**) Time course of DNA relaxation. EcTopA or CtTopA (25 nM) was incubated with 0.3 μg negatively supercoiled DNA for different times (1-30 min). (**d**) Quantification of DNA relaxation based on time course studies. The percent of relaxation was determined by dividing the distance between the negatively supercoiled band (SC); and the weighted center of the partially relaxed band (PR); by the distance between the supercoiled band (SC); and the fully relaxed band (FR). (Formula: percent relaxation = (SC-PR)/(SC-FR)*100 (60). The values are reported as mean ± standard derivation (SD) of results obtained from three independent experiments (also see Figure S4). Statistical comparison between EcTopA and CtTopA was analyzed by Two-Way ANOVA. **P<0.01, *** P<0.001, ****P<0.0001.

**Figure 3.**
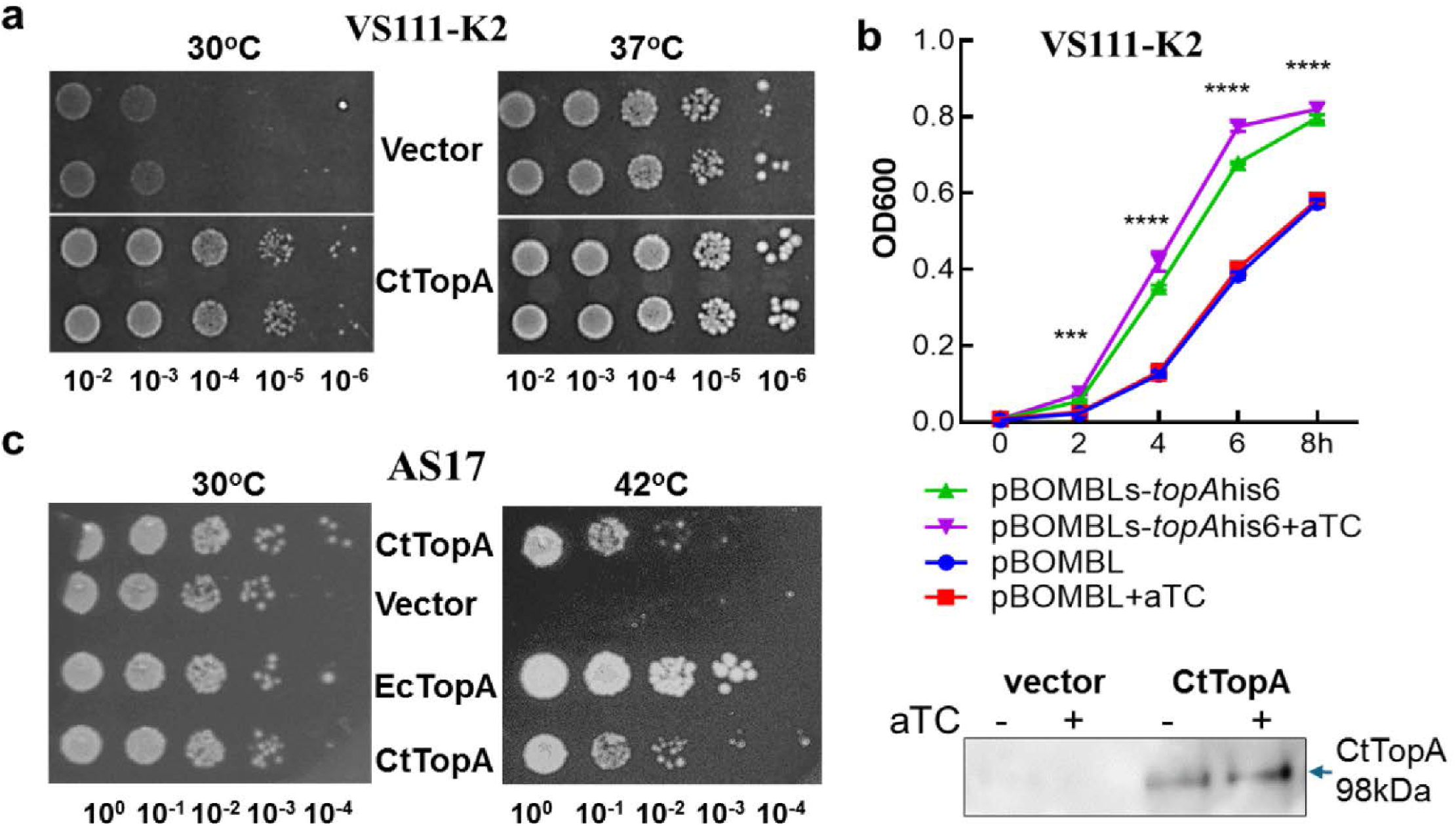
Complementation assay in *E. coli topA* mutant strains. (**a-b**) Results with VS111-K2 transformed with pBOMLs-*topA*His6 expressing CtTopA or empty vector pBOMBLs. Ten-fold serial dilutions of the bacterial cultures were spotted on LB agar plates containing chloramphenicol and spectinomycin. Images were taken 18 h after incubation at 30°C or 37 °C (**a**). Growth curve of *E. coli* strains as indicated at 37°C during 8 h incubation in the presence or absence of aTC at 200 μg/mL(**b**). Y-axis:OD600, x-axis: hours of incubation. Data are presented as mean ± SEM. Statistical comparisons of OD600 between induced and uninduced samples of the same strain were performed by Two-Way ANOVA. ***P<0.001, ****P<0.0001. Lower panel: immunoblotting showing CtTopA expression in *E. coli* with anti-His antibody. Note: leaky expression of CtTopA in the absence of aTC. (**c**) Results with AS17 transformed with plasmid expressing EcTopA or CtTopA as indicated. Ten-fold serial dilutions of the cultures of the transformants were spotted on LB agar plates with kanamycin and incubated at 30°C or 42°C. Images were taken after 18 h for 42°C incubation and 36 h for 30°C incubation. For all strains, two different isolates of *E. coli* transformants were used as biological replicates.

### Characterization of an antibody targeting CtTopA

To better study the domains of CtTopA and to develop novel resources, we produced polyclonal antibodies using two different strategies (Fig. 4a). First, recombinant full-length CtTopA was used as the source of antigen to immunize mice, resulting in anti-CtTopA. Second, we designed and used synthesized peptides containing CtTopA amino acids 737-756 and 843-857 to co-immunize rabbits, resulting in anti-CtTopA_CTD_. By Western blot, we observed that anti-CtTopA and anti-CtTopA_CTD_ specifically recognize an antigen corresponding to ∼98kDa recombinant CtTopA but not recombinant EcTopA or MtTopA (Fig. 4b). Therefore, at least one epitope recognized by both anti-CtTopA and anti-CtTopA_CTD_ has to be situated on the CTD of CtTopA that is not present in EcTopA and MtTopA. These results are in line with the sequence alignment (Figs. 1 and S1) showing differences in the CTD between CtTopA, EcTopA, and MtTopA.

**Figure 4.**
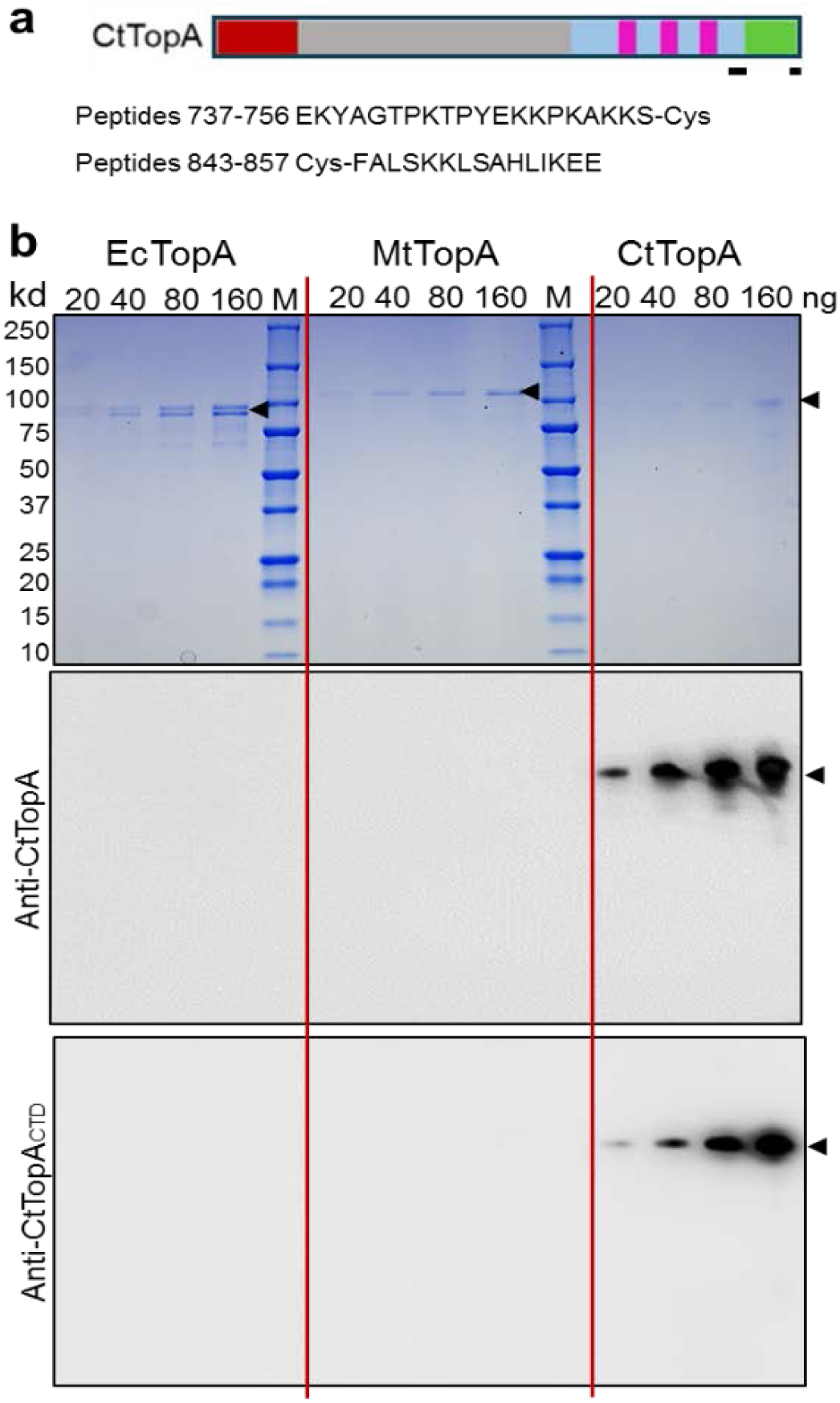
Reaction of anti-CtTopA or anti-CtTopA_CTD_ with the purified recombinant CtTopA. (**a**) Depiction of the scheme for the antigen sources (either full length CtTopA or synthesized peptides) used for antibody production. The location and sequence of peptides are shown. (**b**) Serial dilutions of recombinant EcTopA, MtTopA and CtTopA proteins on SDS-PAGE/coomassie stained gel (upper panel) and immunoblots showing their reactions to anti-CtTopA (middle panel) or anti-CtTopA_CTD_ (lower panel). Arrows show protein bands of interest.

### *C. trachomatis* naturally produces functional SWIB domain-containing CtTopA during infection

To determine whether endogenous CtTopA can be recognized by anti-CtTopA or anti-CtTopA_CTD_ *is situ*, we performed an indirect immunofluorescence assay (IFA). HeLa cells were infected with *C. trachomatis* strains, L2/Nt expressing P*_tet_*-controlled dCas12 and lacking any crRNA and L2/*topA*-kd harboring a CRISPRi plasmid with P*_tet_*-controlled dCas12 and *topA*-specific crRNA; that permitted specific repression of *topA* (20). No signal was detected with anti-CtTopA_CTD_ (data not shown), while anti-CtTopA labeled *C. trachomatis* organisms within the L2/Nt inclusions (Fig. 5a, upper panels). In contrast, only weaker signal was detected in L2/*topA*-kd inclusions, and such signal was further reduced by adding aTC (Fig. 5a, lower panels), consistent with CRISPRi-mediated repression of *topA* expression.

**Figure 5.**
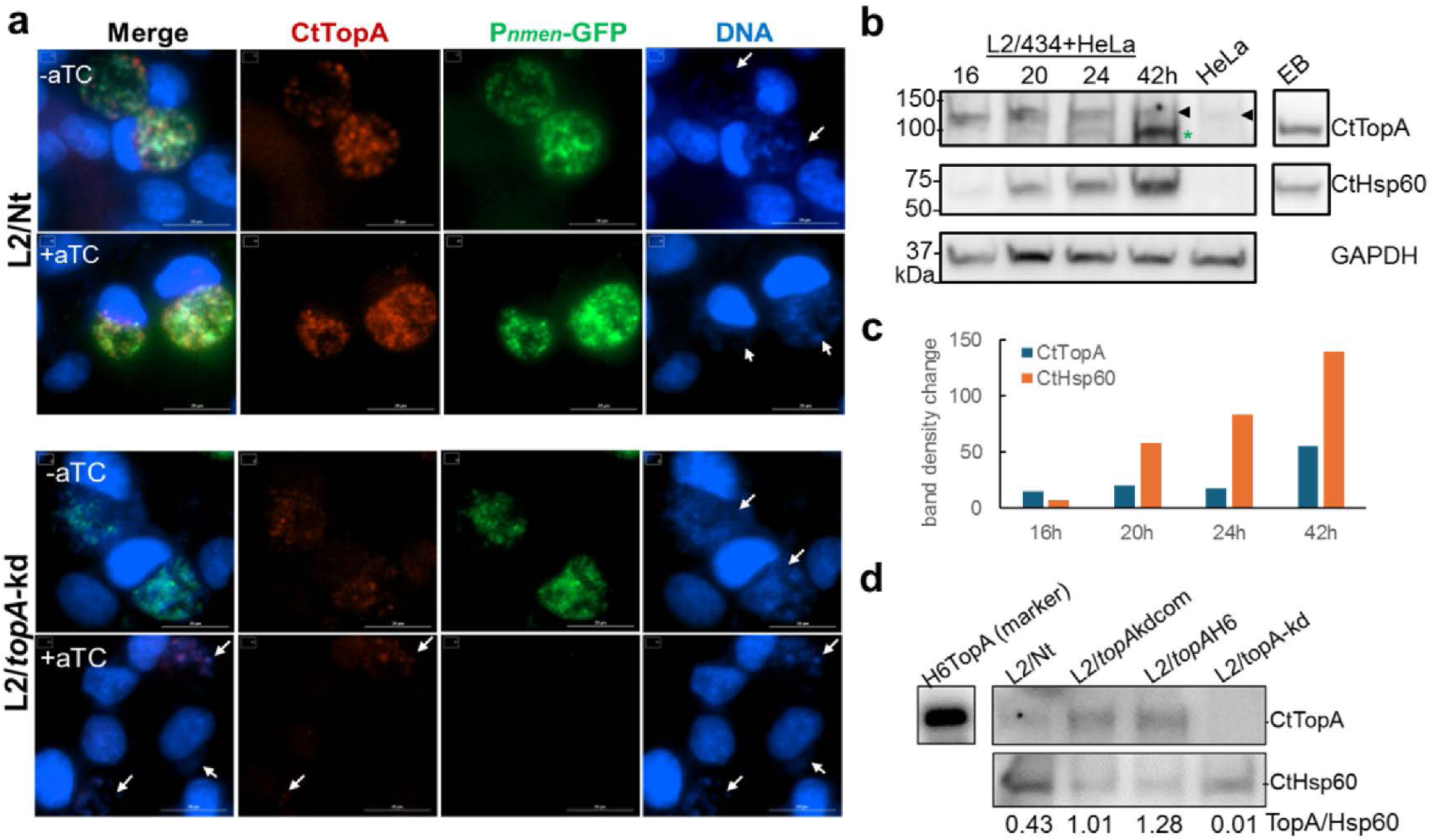
*C. trachomatis* naturally produces SWIB-containing CtTopA. (**a**) Immunofluorescence micrographs of HeLa cells infected with L2/Nt or L2/*topA*-kd at 45 hpi. GFP-expressing chlamydial organisms (green) were stained for CtTopA (red; anti-CtTopA antibody). Cellular and bacterial DNA was counterstained with DAPI (blue). Arrows indicate the location of chlamydial inclusions. Left panels show merged images. Image adjustments of *C. trachomatis* and DNA were applied equally for both bacterial strains and cells. Scale bars=20 μm. (**b**)-(**c**) Immunoblotting of endogenous chlamydial Hsp60 and CtTopA levels in lysates of infected HeLa cells sampled at 16, 20, 24, and 42 h pi. GAPDH was used as a loading control. *: band corresponding to ∼98kDa CtTopA. Arrow: a larger band. Densitometry of the protein band of interest was assessed using ImageJ and presented in (**c**). The full-length blots with the same results were shown in Fig. S3. (**d**) Immunoblotting of CtTopA and CtHsp60 in cell infected with different *C. trachomatis* strains as indicated. Lysates of cells cultured in aTC-containing medium for 40h (4-44 hpi) were used. Values are presented as the density of the CtTopA band normalized to the CtHsp60 band from the same sample using ImageJ.

An immunoblotting analysis of whole cellular lysate was performed to examine the expression pattern of CtTopA protein during *C. trachomatis* infection. The levels of CtTopA at 16-42 h pi were assessed, as *topA* transcripts were predominantly detected at the late stage (17). We observed appearance of CtTopA in L2/434/Bu (nontransformed WT strain) at 20 hpi and later time points (Figs. 5b-c and S3). Similar results were obtained using L2/Nt (data not shown). There were two immunoreactive bands in size around100kDa: one corresponding to ∼98kDa CtTopA and the other one at a larger size. A faint band similar to the larger size was detected in noninfected HeLa cells and only a single band corresponding to ∼98kDa was observed in purified EBs. Thus, the larger band likely represents non-specific binding to a host cell component, which seemed to be induced by *C. trachomatis* infection. The density of ∼98kDa band was hardly detectable when *C. trachomatis* was exposed to chloramphenicol (an inhibitor of bacterial protein synthesis) (Fig. S4), further indicating that it was derived from *C. trachomatis*.

With immunoblotting, we next examined CtTopA expression in *C. trachomatis* strains with *topA* knocked down, complemented, or overexpressed. At 44h pi (the late stage), the CtTopA was detected in L2/Nt, but was faintly detected in L2/*topA*-kd (Fig. 5d), in agreement with the IFA data (Fig. 2a). In contrast, when overexpressing TopA-His6 either in the CRISPRi complemented strain L2/*topA*-kdcom or the L2/*topA*H6 strain lacking CRISPRi elements (20), a band corresponding in size to 98kDa CtTopA was readily detected (Fig.5d). These results indicate *C. trachomatis* expresses CtTopA at mid and late stages and that both endogenous CtTopA and CtTopA overexpressed from a plasmid (in L2/*topA*-kdcom and the L2/*topA*H6) are recognized by anti-CtTopA. It also demonstrates that CRISPRi can effectively repress *topA* transcription and that it, in turn, reduces CtTopA protein levels in *C. trachomatis*.

### Repression of *topA* in *C. trachomatis* inhibits transcripts linked to nucleotide metabolism

We sought to gain further insight into the impact of CtTopA on chlamydial physiology by focusing on nucleotide metabolism. Unlike axenic bacteria, *C. trachomatis* is a nucleotide parasite and relies on its host cells for most of its energy resources during intracellular growth. *C. trachomatis* utilizes two nucleotide transporters to siphon nucleotides from its host cells (33, 34): Npt1 to transport nicotinamide adenine dinucleotide (NAD) and ATP/ADP and the Npt2 to transport GTP, UTP, CTP, and ATP. These transporters presumably serve to compensate for the deficiency in biosynthesis of these molecules *de novo* in *C. trachomatis* except for CTP (35). *C. trachomatis* has a functional CTP synthetase permitting conversion of UTP to CTP in addition to importing host CTP. Repression of *topA* in strain L2/*topA-kd* resulted in growth retardation (Fig. 6a-b), consistent with our previous observations (20). Using RT-qPCR, we assessed the transcription of *C. trachomatis* chromosomal *npt1* and *npt2* genes in strains L2/Nt, L2/*topA*-kd, and L2/*topA*-kdcom. Fig. 6c shows decreases in transcripts of *npt1*, but not *npt2*, at 15 h pi following repression of *topA* in L2/*topA-kd.* This is unsurprising as *npt1* is detectable from the early to the late stages and *npt2* is expressed mainly at middle and late stages (13). The transcript levels of both *npt1* and *npt2* were significantly decreased as compared to the control conditions at 24h pi. These deficiencies could be restored by genetic complementation (in strain L2/*topA*-kdcom) to the levels of the L2/Nt. In contrast, transcription of *pyrG* gene encoding CTP synthetase was not decreased after *topA* was repressed (Fig. S5). The SWIB-containing CtTopA is not conserved beyond the *Chlamydiaceae* family, thus, this may represent a unique function of CtTopA in facilitating nucleotide acquisition from the host cells.

**Figure 6.**
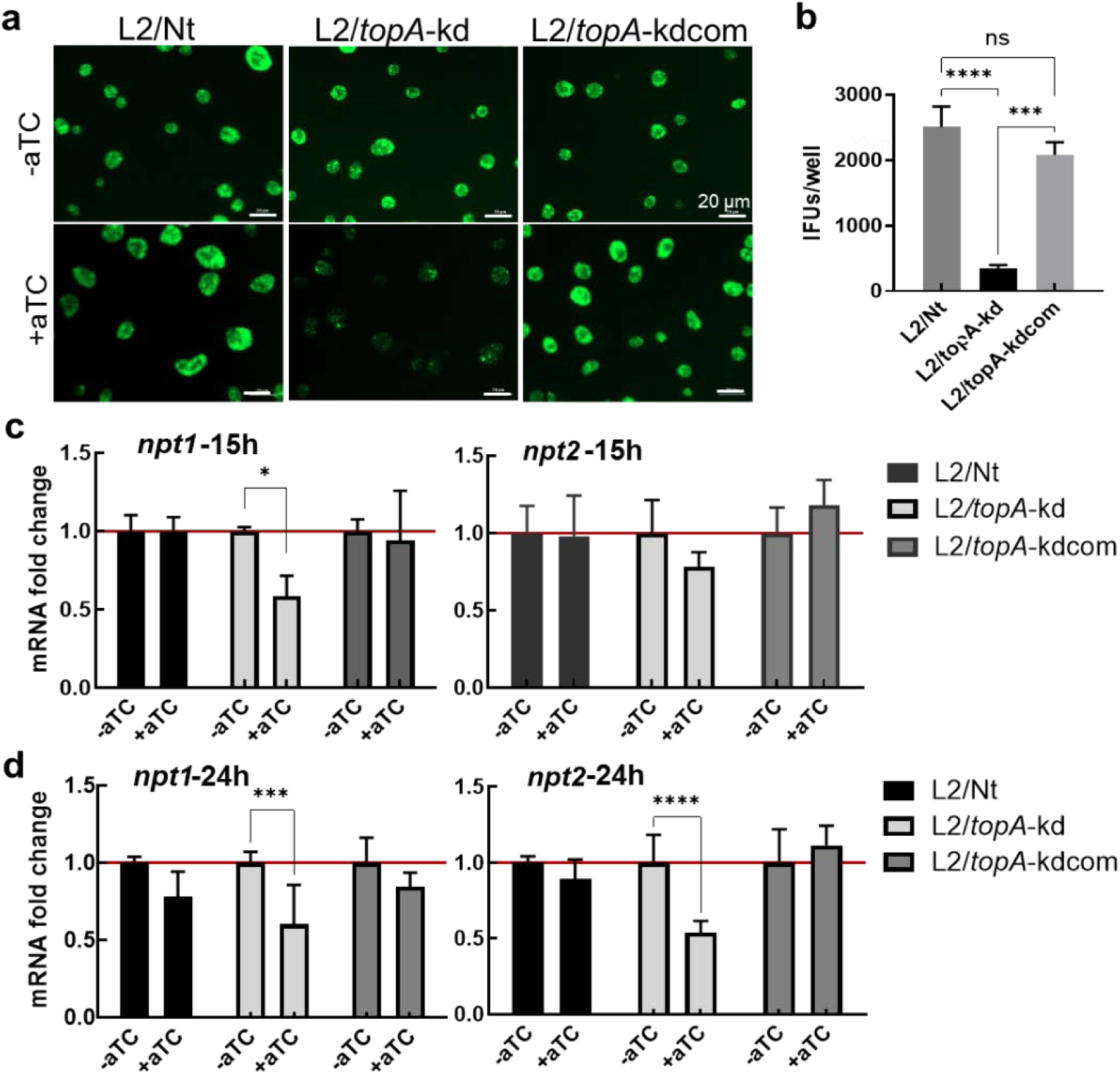
Repression of *topA* induces growth retardation and the decrease in transcription of *npt1* and *npt2*. (**a**) Live-cell images of *C. trachomatis* infected HeLa cells. *C. trachomatis* L2/Nt, L2/*topA*-kd, or L2/*topA*-kdcom at a multiplicity of infection of ∼0.4 were used for infection. Cells were cultured in the absence (-aTC) or presence (+aTC) of aTC from 4 to 24 hpi. Automated imaging acquisition was performed at 24 hpi under the same exposure conditions with Cytation 1. Scale bar = 20 µm. (**b**) Numeration of EB yield using infection assay. The values are presented as mean ± SD from two independent experiments each with three technique repeats. (**c**)-(**d**) Fold change in *npt1* or *npt2* transcript levels. RT-qPCR was conducted with *C. trachomatis*-infected cells grown under inducing (+aTC) or mock inducing (−aTC) conditions starting from 4 hpi for 11 h (to 15 hpi) (c) and 20 h (to 24 hpi) (d). Quantified gene-specific transcripts were normalized to the gDNA levels as determined by qPCR with the same primer pair. The data are presented as the ratio of relative transcript in the presence of aTC to that in the absence of aTC, which is set at 1 as shown by a red line. The values are presented as mean ± SD of two independent experiments each with triplicates. For all panels, statistical significance was determined by One-way or Two-Way ANOVA. ***P ≤ 0.001, ****P ≤ 0.0001.

## Discussion

In the current study, we demonstrate that *C. trachomatis* naturally produces a unique SWIB-containing TopA ortholog. We further show that CtTopA can catalyze DNA relaxation *in vitro*, complement a *topA* mutation in *E. coli*, and is critical for *C. trachomatis* development during infection in HeLa cells. Finally, our data suggest that CtTopA is critical for *C. trachomatis* intracellular growth due, in part, to its ability to regulate genes important for nucleotide acquisition from the host cells. These data indicate that CtTopA exerts its role as a critical virulence factor in chlamydial pathogenesis by facilitating gene regulation and nucleotide metabolism.

Our work demonstrates that CtTopA is distinct from other characterized TopA orthologs in that it has a eukaryotic SWIB-domain at its C-terminus that appears to be conserved in *Chlamydiae* spp. This is unsurprising as *Chlamydia* is an obligate intracellular bacterial parasite that has adapted to survive within eukaryotic host cells and has acquired major *Chlamydia*-specific orthologues with phylogenetic signatures implying eukaryotic origin (15, 20). According to AlphaFold analysis, the predicted protein structure formed by the amino acid sequence of the SWIB domain in CtTopA is likely to have a unique three-dimensional shape that is significantly different from the previously observed structures of other TopA proteins, indicating a potentially novel functional variation within the TopA family. *In vitro* data appear to support this. Our results indicate that the DNA relaxation activity of CtTopA is lower than that of *E. coli* TopA, and CtTopA was unable to complement *E. coli topA* mutants as effectively as the *E. coli* ortholog. This weakened ability could have several explanations. First, the C-terminal zinc ribbon-like domains (D8 and D9) of EcTopA (23) are not found in CtTopA, which instead has the SWIB domain at the C-terminus (Fig. 1). The D8 and D9 of EcTopA have been shown to bind to ssDNA with high affinity (36) and have been proposed to play a significant role in the relaxation activity of EcTopA (37). The relaxation activity of CtTopA would be less efficient if the SWIB domain of CtTopA does not bind the ssDNA region of negatively supercoiled DNA with affinity similar to these zinc ribbon-like domains. Second, the C-terminal zinc finger and zinc ribbon domains of EcTopA also participate in specific protein-protein interactions between *E. coli* TopA and the RNA polymerase (RNAP) (38). Because of the possible lack of or reduced protein-protein interaction with RNA polymerase and its associated proteins, CtTopA may not be as effective as EcTopA in removing transcription-driven negative supercoils during transcription elongation and preventing R-loop formation (39–42), thus limiting the degree of complementation of *topA* mutation in *E. coli*. Future work will look to characterize the protein-protein interactions of full-length or SWIB deleted isoforms of CtTopA in *Chlamydia*. Finally, the degree of complementation by the CtTopA clone in *E. coli* may also be influenced by the plasmid copy number variation (43).

Our data further support that TopA is required for *C. trachomatis* to grow intracellularly. The endogenous TopA protein was expressed at a relatively high level during the middle and late developmental stages of *C. trachomatis*, indicating its temporal action in regulation of developmentally expressed genes. Previously, we reported that expression of late developmental genes (e.g., *hctB* and *omcB*) of *C. trachomatis* was downregulated following *topA* repression while early genes (e.g., *euo* and *incD*) maintained their expression (18). Our observation of a decrease in the transcript levels of the NTP transporters *npt1* and *npt2* suggests that the capacity for nucleotide acquisition from the host cell is reduced when *topA* is repressed and/or that the regulation of these genes is supercoiling sensitive. Such impact is not limited to EB formation, and RB replication may also be affected. In supportive to this, *C. trachomatis* expresses Npt1 and Npt2 in their EB, RB and inclusion membranes as reported previously (44, 45) Although *C. trachomatis* depends upon its host eukaryotic cell for a supply of ATP, GTP, and UTP, it is not auxotrophic for CTP, which can be both transported from the host and synthesized *de novo* by the chlamydial CTP synthetase, PyrG (35, 44). The unchanged transcript levels of *pyrG* observed imply that its transcription is insensitive to reduced DNA relaxation. Under conditions of *topA* repression, *C. trachomatis* likely still gains energy resources for viability via alternative mechanisms, for example, by remodeling the metabolism of host cell mitochondria (46, 47) and hijacking energy metabolites. Because SWIB-containing proteins are not found in prokaryotes aside from *Chlamydia* spp., these results imply a significant aspect of SWIB-containing CtTopA in facilitating the energy acquisition of *C. trachomatis*.

Interestingly, the SWIB-domain has also been predicted in CTL0720/CT460 a 9.7kDa hypothetical protein in *C. trachomatis*. A homolog of CTL0720, Wcw_0377, in *Chlamydia*-like *Waddlia chondrophila* was shown to bind to genomic DNA and to localize in the nucleus when it was expressed in transfected 293T cells (48). Using a CyaA fusion assay, McCaslin et al. (49) detected secretion of CTL0720 in the host cell likely via a type III secretion system (T3SS). We did not observe any co-localization of CtTopA with the nucleus or the cytosol of HeLa cells infected with Ct using IFA, but this negative result could be due to an antibody sensitivity problem or transient translocation of the protein at only specific times during the developmental cycle. Whereas most T3SS effectors utilize an N-terminal signal for secretion, some effectors require a C-terminal signal for proper targeting and interaction with the host cell (50). However, we were unable to identify any such signals in CTL0720 or CtTopA. It is possible that SWIB’s action is context-dependent. For example, there may be overlapping and unique functions of SWIB domains when expressed as a fusion with CtTopA or alone as in CTL0720. In eukaryotes, the SWIB and the MDM2 domains are homologous and share a common fold (51). The SWIB/MDM2 domain superfamily of proteins have diverse functions, including chromatin remodeling (52), p53 regulation (51, 53), and stress response (54) in eukaryotes. Further investigation of the functions of the SWIB-containing proteins, CtTopA and CTL0720, is warranted to understand their role in the *C. trachomatis* developmental program and host-pathogen interactions.

Additional questions remain unanswered. For example, how do the integrated activities of TopA and DNA gyrase contribute to the metabolism of *C. trachomatis*? This is an important question because Topos are drug targets for the development of new antibacterial therapies (4, 55). Recently, Rockey *et al* (56) demonstrated that treatment of cultured *C. trachomatis* with the quinolone ofloxacin at a lethal concentration (1-10 µg/mL) for 72h resulted in metabolic dormancy of *C. trachomatis*. The bacteria could return to active growth after the drug was removed, suggesting the plasticity and sensitivity of *C. trachomatis* in response to Mox-induced DNA relaxation. Although resistance to quinolones is rarely reported in clinical *C. trachomatis* isolates, there are reports showing the potential of acquiring quinolone resistance *via* mutations in the *gyrA* gene after prolonged exposure to sublethal Mox concentrations in culture (57). Mutations in *ygeD*, encoding a putative efflux protein, was also associated with quinolone resistance in clinical isolates (58). Future studies will attempt to evaluate the changes in DNA topology *in C. trachomatis* when altering TopA activity as well as the contributions of gyrase in DNA supercoiling during the chlamydial developmental cycle.

## Materials and Methods

### Reagents

Oligonucleotides and primers were synthesized by Integrated DNA Technologies (Coralville, IA). Restriction enzymes, T4 DNA ligase, and rRNasin were purchased from New England Biolabs (Ipswich, MA). Antibiotics, nucleoside triphosphates, and deoxynucleotide were purchased from ThermoFisher Scientific (Waltham, MA).

### Bioinformatics analysis

The amino acid sequence of CtTopA (CLT0011) and its counterparts in *E. coli*, *M. tuberculosis*, *H. pylori*, *P. aeruginosa*, and *N. gonorrhoeae* were obtained from the UniProt Knowledgebase (UniProtKB) (www.uniprot.org). ClustalW multiple sequence alignment was conducted with Matric BLOSUM62. Domains of CtTopA and its structural model were predicted by InterPro and AlphaFold, respectovely. The amino acid sequence of CtTopA protein was used in protein-protein BLAST (BLASTp) at National Center for Biotechnology Information (NCBI) (https://blast.ncbi.nlm.nih.gov/) searches against the non-redundant standard database corresponding to the *Chlamydiae*/Verrucomicrobia group (taxid:1783257). The selection of 5000 as the maximum number of aligned sequences to display with 0.05 as an E-value-threshold and a BLOSUM62 matrix. NCBI MSA Viewer 1.25.0 was used to visualize amino acid alignment of the SWIB-domain regions from the top 500 homologues of CtTopA.

### Expression of recombinant TopAs in *E. coli*

The strains and plasmids used in this study are listed in Table S2. The coding sequence of CtTopA optimized for expression in *E. coli* was custom synthesized by Gene Universal (Delaware, USA) and inserted into vector pET28a(+) for expression of recombinant CtTopA with N-terminal 6xHis tag. The resulting pET-CtTopA plasmid was transformed into *E. coli* strain BL21(DE3). Transformants were grown in LB (Miller) broth with 50 μg/ml kanamycin at 37°C for overnight culture. Next day, the overnight cultures were diluted 1:100 in LB with 50 μg/ml kanamycin and grown until OD600 reached 0.4. Recombinant protein expression from the T7 promoter was induced with the addition of 1 mM IPTG. The cells were harvested after additional growth at 37°C for 4 hr. Similarly, plasmid expressing EcTopA (37) or MtTopA (59) (Table S2) were used for expression of these recombinant topoisomerases in BL21 STAR (DE3) strain (Invitrogen) and C41(DE3) (Lucigen) respectively.

### Purification of recombinant CtTopA, EcTopA, and MtTopA

EcTopA and MtTopA were purified as previously described (37, 59). For purification of CtTopA, the pelleted bacterial cells were resuspended in buffer of 50 mM sodium phosphate pH 7.4, 0.3 M NaCl, 20 mM imidazole. After addition of 1 mg/mL lysozyme, the cells were left on ice for 1 hr before three cycles of freeze-thaw to lyze the cells. The soluble lysate obtained after centrifugation at 32000 rpm for 2 hr was mixed with Ni-NTA agarose (from Invitrogen, Thermo Fisher) and packed into a column. After washing, the protein was eluted with buffer of 50 mM sodium phosphate pH 7.4, 0.3 M NaCl, 250 mM imidazole. Protein concentration was determined with the Bradford assay.

### *In vitro* assay of topoisomerase relaxation activity

The relaxation activity assay was conducted in 20 μl of 10 mM Tris–HCl, pH 8.0, 50 mM NaCl, 0.1 mg/ml gelatin, 2 mM MgCl_2_ wiith 0.3 μg of negatively supercoiled pBAD/thio plasmid DNA as substrate. Following addition of topoisomerase, the reactions were incubated at 37°C for the length of time indicated in the results and stopped by the addition of 4 μl of stop solution (50 mM EDTA, 50% glycerol and 0.5% v/v bromophenol blue). The supercoiled DNA substrate and relaxed DNA products were separated by electrophoresis in a 1% agarose gel with TAE (40 mM Tris-acetate, pH 8.0, 2 mM EDTA) buffer. Following staining with 1 μg/ml ethidium bromide, the gel was de-stained with deionized water and then photographed with UV light and the Alpha Imager Mini. Percent relaxation was determined as previously described (60). The migration distance of supercoiled (SC) DNA, fully relaxed (FR) DNA and partially relaxed (PR) DNA bands were identified using AlphaViewer. The weighted distance of PR bands for each lane was calculated from the data obtained. The % relaxation was calculated with the formula (SC-PR)/(SC-FR) *100.

### Complementation of *topA* mutations in *E. coli*

The *E. coli*-*C. trachomatis* shuttle plasmid pBOMBLs-*topA*H or vector control pBOMBLs were transformed into *E. coli* strain VS111-K2 (Table S2). The resulting transformants were grown in LB (Miller) broth with 50 μg/ml spectinomycin at 37°C overnight. The cultures were diluted to OD600 = 0.1, prior to 10 fold dilution and then dropped on LB agar. Plates were incubated at 30, 37, and 42°C and imaged at 18 h. For growthe curve, the overnight culture was diluted in fresh medium at a ratio of 1:100 and cultured in LB medium with or without aTC at 37°C. Culture were sampled to measure the optical density at 600 nm (OD600) every 2 hrs. The absorbance values were plotted against the growth time.

The pET-CtTopA plasmid was transformed into the *E. coli* AS17 (Table S2), which has a temperature-sensitive *topA* mutation and requires complementation for growth at 42°C (32, 59). The LIC-ETOP plasmid (37) expressing His-tagged *E. coli* TopA from the T7 promoter was used as positive control for comparison along with empty vector as negative control. Individual transformants with pET-CtTopA were first isolated at 30°C as biological replicates. The AS17 transformants were grown in LB (Miller) broth with 50 μg/ml kanamycin at 30°C overnight to saturation. The cultures were first diluted with LB for OD600 value to equal 0.1 before ten-fold serial dilutions were prepared for spotting of 5 μl of each dilution onto LB agar plates with 50 μg/ml kanamycin. The plates were photographed following incubation at 30°C for 36 hr or 42°C for 18 hr.

### Cell culture and *C. trachomatis* infection

HeLa 229 cells (human cervical epithelial carcinoma cells; ATCC CCL-2) were cultured in RPMI 1640 medium (Gibco) containing 5% heat-inactivated fetal bovine serum (Sigma-Aldrich), gentamicin 20 µg/mL, and L-glutamine (2 mM) (RPMI 1640–10) at 37°C in an incubator with 5% CO_2_. Cells were confirmed to be *Mycoplasma* negative by PCR as described previously (61). To propagate and prepare the large amounts of EBs, HeLa cells grown in T175 flasks were infected with *C. trachomatis* (Table S2) and cultured in RPMI 1640–10 at 37°C for 45 h pi. For transformed strains, the medium was supplied with spectinomycin (500 µg/mL). Cells were harvested for EB purification as described previously (34). The purified EB pellet was resuspended in sucrose-phosphate-glutamic acid buffer (10 mM sodium phosphate, 220 mM sucrose, 0.50 mM l-glutamic acid). The EB aliquots were stored at −80°C until use. Serial dilutions of EBs were used to determine the titers in 96-well plates as inclusion-forming units (IFU). For phenotypic analysis, *C. trachomatis* EBs was used to infect cells grown in 96-well plates (catalog #655090, Greiner) with a dose that results in ∼30% to 40% of cells being infected. After centrifugation with a Beckman Coulter model Allegra X-12R centrifuge at 1,600 × *g* for 45 min at 37°C, fresh medium was added to the infected cells and incubated at 37°C for various time periods as indicated in each experimental result. For comparison, different strains were infected side-by-side in the same culture plate with a setup of at least triplicate wells per condition.

### CtTopA antibody production

Purified full-length His6-CtTopA was used to produce polyclonal antibody against chlamydial TopA in mice as described previously (62). Briefly, 50 ug of recombinant TopA emulsified in equal volumes of complete Freund’s Adjuvant were intraperitoneally injected into a mouse. Two weeks later, the same amount of TopA antigen, emulsified in incomplete Freund’s Adjuvant, was similarly injected twice at an interval of two weeks. Sera were collected two weeks after the final booster injection. The synthesized peptides corresponding to amino acids 737-756 and 843-857 of CtTopA were used to produce antibody in rabbit (Pacific Immunology). The final serum was purified by an affinity column.

### Immunofluorescence assay (IFA) and image analysis

For IFA, *C. trachomatis*-infected HeLa cells cultured for 42 hpi were fixed with 4% formaldehyde for 15 min and permeabilized by using 0.1% Triton X-100 for 15 min, followed by immunostaining with mouse anti-CtTopA (1:800) or rabbit anti-CtTopA_CTD_ (1:500) overnight at 4°C. After extensively washing, cells were then incubated with Alexa Fluor 568-conjugated secondary antibody (1:200; Molecular Probes) for 45 min at 37°C and counterstained with DAPI (4′,6-diamidino-2-phenylindole dihydrochloride). Images were automatically captured at ×20 magnification using the Cytation 1 multimode reader (BioTek Instruments, Winooski, VT), followed by processing and analyzing with Gen5 software.

### *C. trachomatis* enumeration and end point one-step growth curve

IFU assays were performed in 96-well plates to determine yield of EB progeny. *C. trachomatis*-infected cells in culture plates were frozen at −80°C, thawed once, scraped into the medium, serially diluted, and then used to infect a fresh monolayer of HeLa cells. The infected cells were cultured in RPMI 1640–10 with 500-µg/mL spectinomycin at 37°C for 40 h. Cells were then fixed with 4% paraformaldehyde, permeablized with 0.1% triton X-100, and stained with mouse monoclonal antibody against the major outer membrane protein (MOMP) of *C. trachomatis* LGV L2 (31). Images were taken using fluorescence microscopy, and the inclusion numbers in triplicate wells were counted.

### Immunoblotting

*C. trachomatis*-infected cells in 12-well culture plate were lysed directly in 8 M urea buffer containing 10 mM Tris-HCl (pH 8.0), 0.1% SDS, and 2.5% β-mercaptoethanol. The protein content was determined by a bicinchoninic acid protein (BCA) assay kit (Thermal Fisher). Cellular lysate was prepared from each samples and an equal amount of protein was loaded into a single lane of the 4-15% SDS-polyacrylamide gel (BioRad). After electrophoresis and transfer to a polyvinylidene difluoride (PVDF) membrane (Millipore), the membrane was individually incubated with antibody specific to CtTopA (1:1000), Hsp60 (1:500) (63), or the loading control host GAPDH (1:2000) (MilliporeSigma), followed by incubation with the HRP-conjugated secondary antibody. The blot was imaged on an Azure c600 imaging system. The relative density of a given protein band is evaluated across its respective row by ImageJ.

### DNA and RNA analysis

DNA and RNA were simultaneously extracted from *C. trachomatis*-infected HeLa cells using the Quick-DNA/RNA miniprep kit (Zymo), and their concentrations were determined using a NanoDrop spectrophotometer (Thermo Scientific). For real-time RT-qPCR, a total of 2 μg of RNA was reverse transcribed into cDNA using a high-capacity cDNA reverse transcriptase kit according to the manufacturer’s instruction (Thermo Fisher). Dilutions of cDNA were then used for amplification of the genes of interest in total volumes of 20 μL with appropriate primers (see below) using PowerUp SYBR green master mix (Thermo Fisher). For real-time qPCR analysis, DNA samples were used as the templates to amplify genes of interest in 20-μL reaction mixture volumes. Each sample was run in triplicate in a 96-well plate on a real-time PCR system (Bio-Rad). The following conditions were used: 95°C for 3 min, and then 95°C for 5 s and 63°C for 30 s. The last two steps were repeated for 40 cycles, with fluorescence levels detected at the end of each cycle. The quantifications of qPCR or RT-qPCR products were calculated from the standard curves with chlamydial genomic DNA from purified EBs as templates. To amply chlamydial *npt1*, *npt2*, and *pyrG*, the following primer pairs were used individually: rt_*npt1*F/rt_*npt1*R (5’-TTGGCCGATACACATGCATG-3’)/(5’-TCCCGGTGCTGTAACGATAA-3’), rt_*npt2*F/rt_*npt2*R (5’-TCCCTATGGCCGTAGATCCT-3’)/(5’-ACGTGTCATCCATCAGCGA-3’), and CT183*pyrG*_F (5’-AAGTATACGTGACCGACGATG-3’)/ CT183*pyrG*_R (5’-CTGCGCACGATTGAATGACAT-3’).

### Statistical analyses

Data analyses were performed using Prism (version 10; GraphPad, San Diego, CA). Statistical significance was determined by one-way or two-way analysis of variance (ANOVA) as indicated in each result. P values of <0.05 were considered statistically significant.

## ACKNOWLEDGMENTS

Research reported in this publication was supported in part by National Institutes of Allergy and Infectious Diseases grants R21AI175651 to both L.S. and S.P.O and R35GM139817 to Y.T. Caitlynn Diggs is a scholar of NIH Postbaccalaureate Research Education Program (PREP) (1R25GM148309).

